# NKG7 is a stable marker of cytotoxicity across immune contexts and within the tumor microenvironment

**DOI:** 10.1101/2025.02.05.636622

**Authors:** Roberta Turiello, Susanna S. Ng, Elisabeth Tan, Gemma van der Voort, Nazhifah Salim, Michelle Yong, Malika Khassenova, Johannes Oldenburg, Heiko Rühl, Jan Hasenaur, Laura Surace, Marieta Toma, Tobias Bald, Michael Hölzel, Dillon Corvino

**Author notes:** Contact info: Correspondence. authors contributed equally (Roberta Turiello, Susanna S. Ng, Michael Hölzel, Dillon Corvino). Lead contact: Further information and requests for resources and reagents should be directed to and will be fulfilled by lead contact, Dillon Corvino.

## Abstract

Cytotoxicity is a cornerstone of immune defense, critical for combating tumors and infections. This process relies on the coordinated action of granzymes and pore-forming proteins, with Granzyme B (GZMB) and Perforin (*PRF1*) being key markers and the most widely studied molecules pertaining to cytotoxicity. However, other human granzymes and cytotoxic components remain underexplored, despite growing evidence of their distinct, context-dependent roles. Natural Killer Cell Granule Protein 7 (NKG7) has recently emerged as a crucial cytotoxicity regulator, yet its expression patterns and function are poorly understood. Using large publicly available single-cell RNA sequencing atlases, we performed a comprehensive profiling of cytotoxicity across immune subsets and tissues. Our analysis highlights NKG7 expression as a strong marker of cytotoxicity, exhibiting a strong correlation with overall cytotoxic activity (r = 0.97) and surpassing traditional markers such as Granzyme B and Perforin in reliability. Furthermore, NKG7 expression is notably consistent across diverse immune subsets and tissues, reinforcing its versatility and robustness as a cytotoxicity marker. These findings position NKG7 as an invaluable tool for evaluating immune responses and a reliable indicator of cytotoxic functionality across biological and clinical contexts.

## Introduction

Immunotherapy has revolutionized cancer treatment, significantly improving patient prognosis and, in some cases, offering curative potential, particularly in hematological malignancies. Most immunotherapy strategies leverage the anti-tumor capabilities of CD8+ T-cells and Natural Killer (NK) cells. These cells exert anti-tumor functions through the release of specific cytokines, death-receptor signaling, or the release of cytotoxic granules. The deployment of cytotoxic granules is especially critical, as they induce targeted, rapid apoptosis of target cells. Cytotoxic granules are specialized secretory lysosomes that release their cytotoxic payload into the immunological synapse, consisting of granzymes, perforin, and granulysin, into the immunological synapse (de Saint Basile et al., 2010).

Granzymes are a core component of the cytotoxic granule-mediated death machinery. These serine proteases cleave various intracellular substrates to initiate target-cell death. In humans, five granzymes (A, B, H, K, and M) have been identified, each with differing substrate specificities and thus, the capacity to induce distinct forms of cell death (Chowdhury & Lieberman, 2008). However, the contexts, heterogeneity and dynamics of granzyme expression remain poorly understood. Originally considered redundant, granzymes are now increasingly recognized for their distinct and specialized functions. For example, granzyme A and B - the most well-studied members of the granzyme family - induce caspase-independent pyroptosis and caspase-dependent apoptosis respectively (Adrain et al., 2005; Beresford et al., 1999; Shresta et al., 1999; Zhou et al., 2020). Meanwhile granzymes H, K, and M remain poorly understood but exhibit unique functions, including the induction of alternative apoptosis pathways, microtubule disruption, cytokine processing, extracellular matrix remodeling, and modulating inflammatory responses (Fellows et al., 2007; Hay & Slansky, 2022). These latter processes highlight some of the non-cytotoxic functionalities increasingly being attributed to granzyme activity. Perforin is a key mediator of cytotoxicity and predominantly functions to facilitate the delivery of cytotoxic effectors such as granzymes and granulysin into target cells (Voskoboinik et al., 2015). Granulysin is involved in cytotoxicity and anti-microbial response, contributing to anti-tumoral responses through membrane disruption and immune modulation (Krensky & Clayberger, 2009). Together these granule components orchestrate the rapid and efficient cytotoxic response while engaging in non-cytotoxic roles, that can be either pro-tumorigenic or anti-inflammatory.

Tumors and pathogens have evolved mechanisms to evade cytotoxicity, including overexpressing specific granzyme inhibitors or downregulating granzyme targets, thus reducing their susceptibility to particular granzymes (Tuomela et al., 2022). This highlights the potential benefits of granzyme heterogeneity. For instance, granzyme B is selectively inhibited by molecules such as Serpin-B9, while the other granzymes are unaffected (Kaiserman & Bird, 2010). Similarly, granzyme H degrades an adenoviral inhibitor of granzyme B (Andrade et al., 2007). This evolutionary interplay underscores the necessity for the diverse yet overlapping functions of the granzyme family.

In recent years, Natural Killer cell Granule Protein 7 (NKG7) has emerged as a potent marker of cytotoxic populations. Increasingly, *NKG7* expression is used to identify cytotoxic populations in sequencing datasets; however, research into the expression and immunological role of NKG7 is in its infancy. Specifically, the function and expression dynamics of NKG7 are largely unknown. NKG7 was first identified to be expressed in the cytotoxic granules of NK and T-cells and has subsequently been shown to regulate anti-tumoral effector functions (Li et al., 2022; Medley et al., 1996; Ng et al., 2020; Turman et al., 1993; Wen et al., 2022). It is thought that NKG7 is involved in the release of cytotoxic granules during the immune response (Lelliott et al., 2022a; Li et al., 2022; Ng et al., 2020). Clinically, NKG7 expression is associated with improved patient outcomes across various tumor entities. (Ayers et al., n.d.; Durante et al., 2020; Fairfax et al., 2020; Li et al., 2022; Szabo et al., 2019) Altogether, current findings demonstrate the importance of NKG7 in regulating anti-tumoral cytotoxicity, and its utility for the assessment and prediction of clinical responses to immunotherapy.

Granzyme functions are diverse, yet much remains unknown about this heterogeneity and it is often underappreciated. The majority of research has focused on granzymes A and B, overlooking the broader diversity of granzyme functions. NKG7, a newly emerging player in cytotoxicity, shows promise as a strong correlate of cytotoxic activity. In our study, we comprehensively profiled cytotoxic molecule usage across cytotoxic and non-cytotoxic populations in healthy patients, various tissues, and disease settings. We found NKG7 expression outperforms traditional markers of cytotoxicity such as granzyme B and perforin, in capturing cytotoxic populations. Furthermore, NKG7 was found to be stably expressed across tissues, cell subsets, and disease conditions. While we confirmed some expected expression patterns such as expression of granzyme B in pDCs, we also identified previously overlooked markers (Jahrsdörfer et al., 2010). For example, a notable proportion of effector cells lack significant perforin expression. Our findings suggest NKG7 may serve as a valuable pan-cytotoxicity marker, crucial for identifying cytotoxic cells despite inherent granzyme heterogeneity.

## Results

### Cytotoxicity profiling reveals conserved and distinct patterns of cytotoxic molecule expression in human PBMCs

To investigate cytotoxic molecule expression, we utilized a scRNAseq dataset of PBMCs from healthy donors (Hao et al., 2021). This dataset, derived from multimodal RNA and protein sequencing, encompasses well-defined cell subsets (Figure 1A). Scoring cells for cytotoxicity (*GZMA*, *GZMB*, *GZMH*, *GZMK*, *GZMM*, *GNLY*, *PRF1*, and *NKG7*) reveals a high density of cytotoxic molecule expression in NK-Dim (CD56DimCD16+) and CD8-EM (Effector Memory) (Figure 1B-C). Inspection of cytotoxicity score reveals NK-Dim, proliferative NK, and CD8 populations as high expressers of cytotoxicity markers (Figure 1C). Despite being classically described as an immature NK cell subset and pro-inflammatory subset (Farag & Caligiuri, 2006), NK-Bright (CD56BrightCD16-) cells scored highly for cytotoxicity, with levels similar to what is seen in classical cytotoxic populations such as CD4-CTLs and CD8-EM cells. Innate-like populations also showed diverse levels of cytotoxicity ranging from highly cytotoxic (gdT-V9D2) to poorly cytotoxic (MAIT) populations. Strikingly, gdT cells exhibited a bimodal cytotoxicity distribution with sub-populations both high and low in cytotoxicity (Hartigan’s dip test, *p* < 2.2e-16). To further probe these results, the expression of individual markers was evaluated across immune subsets (Figure 1D). This interrogation revealed both expected and poorly described or seldom appreciated granzyme expression patterns. For example, the classic cytotoxic subsets (CD8-EM, NK-Dim, and CD4-CTLs) all had similar and expected expression patterns. These subsets expressed high levels of all cytotoxic molecules except *GZM*K, which was low or absent. A similar expression pattern was observed in gdT and proliferating NK subsets. Conversely, NK-Bright cells, which are typically described as pro-inflammatory or regulatory in nature, expressed high levels of all markers except *GZMH*. Interestingly, the gdT-V9D2 population could be distinguished from remaining gdT cells by the expression of *GZMK*. Additionally, gdT-V9D2 along with CD8-Prolif cells were the only two populations assessed that showed appreciable levels of expression of all cytotoxic molecules. Meanwhile, MAIT cells expressed high levels of expression of all cytotoxic molecules except *GZMB* and *GZMH*. This reveals that the low cytotoxicity score observed for MAIT cells is driven by the lack of *GZMB* and *GZMH*. Of note, pDCs displayed high expression of *GZMB* but no other cytotoxic molecule. Additionally, a low level of expression of *GZMA*, *GZMM*, *GNLY*, and *NKG7* was detected in platelets. Evaluation of the cytotoxic landscape of human PBMCs revealed known and unexpected expression patterns for cytotoxic molecules. This analysis reveals that populations not typically considered as cytotoxic can express appreciable levels of cytotoxic molecules. Furthermore, certain immunological subsets appear to prefer the expression of particular cytotoxic molecule patterns.

**Figure 1:**
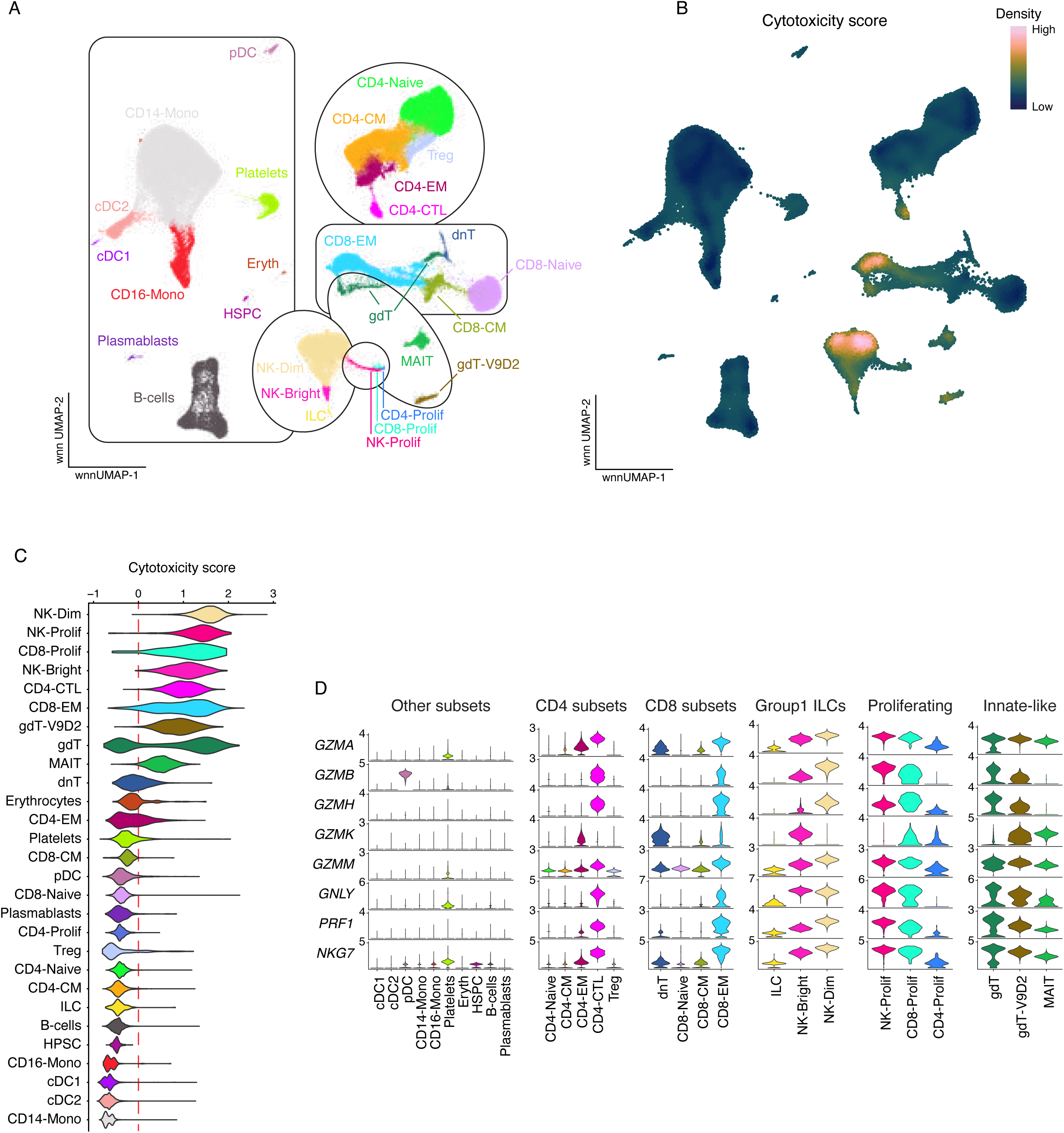
Pattern of cytotoxic molecule expression in human PBMC subsets. (A) The cellular subsets of human PBMC as identified using integrated scRNAseq and scCITEseq data visualized on weighted-nearest neighbor (wnn) UMAP coordinates. (B) Density of expression for cytotoxicity score overlayed on wnnUMAP. (C) Violin plots showing the “cytotoxicity score” across immune populations (D) Violin plots showing the imputed expression of cytotoxicity markers that contribute to the cytotoxicity score. Cellular subsets are abbreviated as follows: CD14⁺ Monocyte (CD14-Mono), CD16⁺ Monocyte (CD16-Mono), CD56BrightCD16⁻ (NK-Bright), CD56DimCD16⁺ (NK-Dim), Central Memory (CM), Conventional Dendritic Cell Type 1 (cDC1), Conventional Dendritic Cell Type 2 (cDC2), Cytotoxic T-lymphocyte (CTL), Double-Negative T-cell (dnT; CD4⁻CD8⁻ T-cells), Effector Memory (EM), Gamma Delta T-cell (gdT), Hematopoietic Stem and Progenitor Cell (HPSC), Innate Lymphoid Cell (ILC), Mucosal-Associated Invariant T-cell (MAIT), Plasmacytoid Dendritic Cell (pDC), Proliferating (Prolif), Regulatory T-cell (Treg), Vγ9Vδ2 Gamma Delta T-cells (gdT-V9D2).

### NKG7 is a reliable marker of cytotoxicity in human PBMC subsets

Given the diverse patterns of cytotoxic molecule expression observed in human PBMCs, we sought to further evaluate the dynamics of cytotoxic molecule use. The correlation of each cytotoxic gene with one another was evaluated within the two most cytotoxic subsets (CD8 and NK cells). This revealed *GZMK* as poorly and inversely correlated with the expression of other cytotoxic genes in both CD8 and NK subsets (Supplementary Figure 1A). This prompted us to ask, which cytotoxic gene is most strongly and consistently correlated with cytotoxicity across all subsets present within human PBMCs. To evaluate this, subsets were iteratively scored for cytotoxicity using a signature of all cytotoxic genes except the gene of interest. The correlation between the gene of interest and overall cytotoxicity score was then determined and visualized. The analysis revealed that *NKG7* expression had the strongest correlation (r = 0.97) with cytotoxicity score (Figure 2A). *PRF1*, *GNLY*, and *GZMA* (r = 0.95, r = 0.95, r = 0.94; respectively) all had similar correlation values with cytotoxicity score. Interestingly, *GZMB* had the second lowest correlation score (r = 0.66), which was driven by the unique and singular expression of *GZMB* in pDC’s. Excluding pDC cells from analysis resulted in a correlation score of r = 0.9 for *GZMB* (Supplementary Figure 1B).

**Figure 2:**
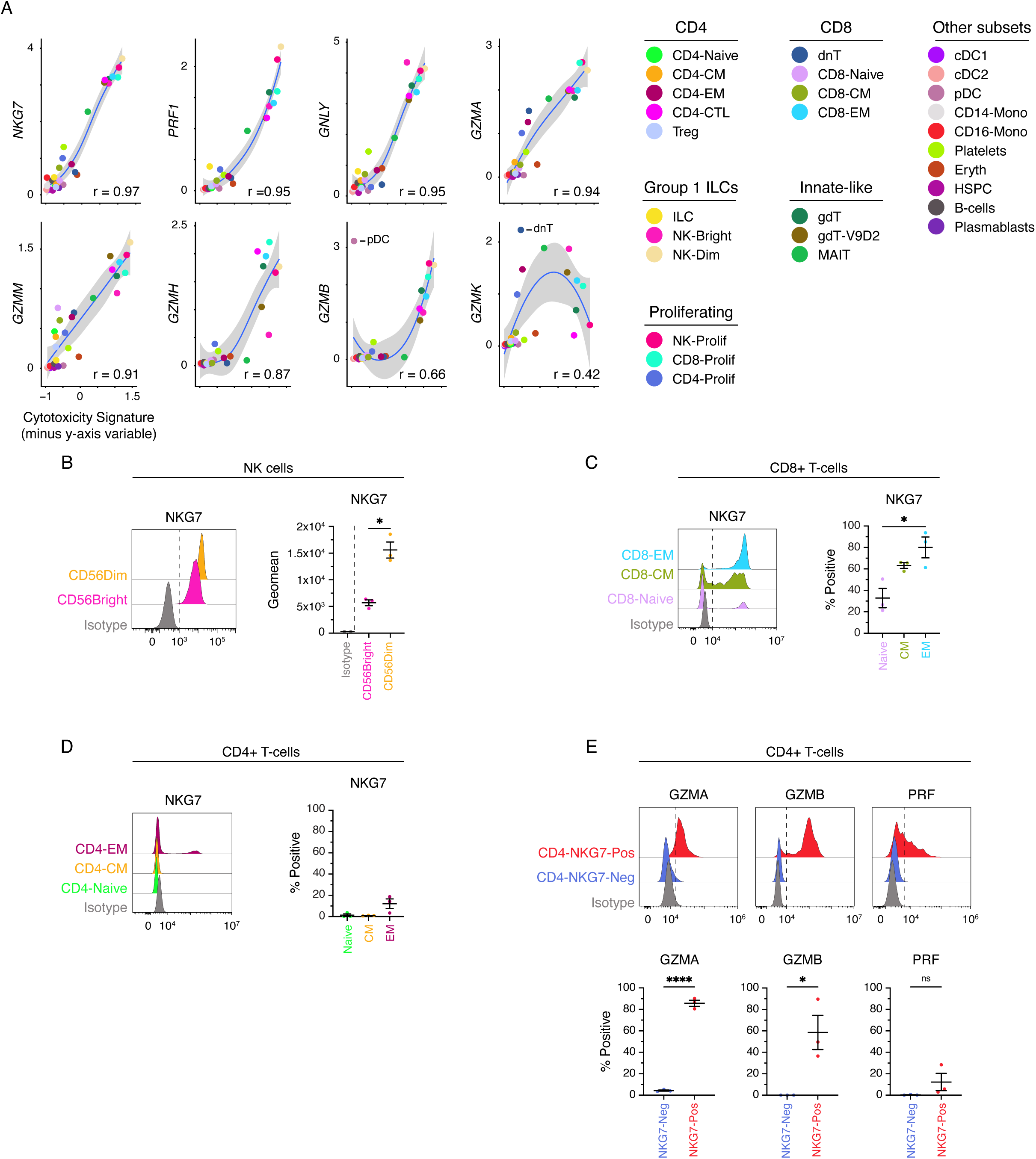
NKG7 correlates with cytotoxicity across human PBMC subsets. (A) Scatterplots demonstrating the correlation between y-axis gene expression and cytotoxicity signature across PBMC subsets. Pearson correlation is displayed and the shaded area represents the 95% CI. (B) Histograms showing the expression of NKG7 in NK cell populations and the corresponding geometric mean fluorescence intensity in CD56Bright and CD56Dim populations. P values from Welch’s t test. (C–D) Histograms showing the expression of NKG7 in EM, CM and Naïve CD8+ T-cells or EM, CM and Naïve CD4+ T-cells, respectively; and dot plots showing the frequency of NKG7+ populations. P values from ordinary one-way ANOVA. (E) Histograms showing the expression of GZMA, GZMB, PRF and in NKG7 positive and negative CD4+ T-cells, and corresponding dot plots indicating the frequency of positive cells. Dashed lines in the histograms do not represent gating thresholds but are included for visual comparison. Gating was determined based on unstained controls, isotype controls, or fluorescence minus one (FMO) controls, depending on the most appropriate approach for each marker. P values from unpaired t-test. * = P < 0.05; **** = P value < 0.0001.

Given these findings, and the known discordance of RNA and protein, we sought to investigate the expression patterns of NKG7 at the protein level. Interestingly, limited studies have evaluated the protein expression of NKG7, perhaps due to limited reagent availability. As such, we utilized the anti-TIA1 antibody clone 2G9A10F5 (herein referred to as 2G9) (Medley et al., 1996). This monoclonal antibody recognizes a pentameric epitope (GYETQ) at the C-terminus of TIA1. Similarly, the C-terminus of NKG7 contains the pentameric GYETL sequence (Supplementary Figure 1C). Others have established 2G9 as a cross-reactive antibody capable of binding both NKG7 and TIA1 (Medley et al., 1996). We validated these findings using transfected HEK293T cells over-expressing tagged TIA-1 or NKG7. Western blot analysis confirmed that 2G9 is cross-reactive for NKG7 and TIA-1 (Supplementary Figure 1D). *NKG7* and *TIA1* genes are encoded on separate chromosomes, with *NKG7* on chromosome 19 and *TIA1* on chromosome 2. Furthermore, these genes have different expression patterns, predicted structures, and functions. Therefore, *TIA1* is not expected at appreciable levels within immunological subsets. To verify this, we evaluated the expression of *NKG7* or *TIA1* in total or sorted PBMC subsets using both bulk and single-cell RNAseq datasets (Supplementary Figure 1E–G). This revealed minimal to no *TIA1* transcript can be detected across immunological subsets. This observation was further validated at the protein level where TIA1 was not detected in PBMC lysates incubated with the anti-TIA-1 antibody EPR9304 (Supplementary Figure 1H). Therefore, it can be assumed that signal from 2G9 in PBMCs derives from NKG7.

Hence, we evaluated the protein expression of NKG7 in cytotoxic subsets from healthy PBMC samples using the 2G9 antibody. This verified NK cells as potent expressers of NKG7 with both CD56Bright and CD56Dim populations expressing NKG7. However, on a per-cell basis, CD56Dim NK cells exhibited significantly higher NKG7 expression levels compared to CD56bright cells (Figure 2B). In line with transcriptomic data, the frequency of NKG7 expression in CD8 T-cells increased with differentiation state (Figure 2C). Similarly, NKG7 protein expression in CD4 T-cells mirrored transcriptomic trends, with the increased levels observed in the CD4-EM subset (Figure 2D). However, this increase did not reach statistical significance, as only a small proportion (∼12%) of CD4-EM cells were positive for NKG7. Interestingly, phenotypic identification of CD4-CTLs is still a matter of debate (Malyshkina et al., 2023; Speiser et al., 2023). However, *NKG7* has appeared in numerous transcriptomic signatures of CD4-CTLs. (Preglej & Ellmeier, 2022). Indeed, we found that NKG7 positive CD4 T-cells were enriched for cytotoxic molecules such as GZMA, GZMB, and PRF (Figure 2E). Therefore, NKG7 may serve as a valuable phenotypic marker to capture CD4-CTLs.

### NKG7 correlates with cytotoxicity across disparate tissues

It was previously observed that *NKG7* strongly correlates with cytotoxicity in healthy PBMC subsets. However, the tissue-specific expression patterns of *NKG7* and other cytotoxic molecule genes are poorly described. To address this, we took the Tabula Sapiens scRNAseq dataset and probed the expression of cytotoxic genes across NK and CD8+ T-cell from various immunologically relevant organs. This revealed that *NKG7* is consistently expressed across NK cells from various tissue locations (Figure 3A). In contrast, genes such as *GZMH* or *GZMK* showed dynamic expression differences between NK cells from different tissues. This possibly reflects known and expected differences in the abundance of NK subsets in these tissues and their distinct expression profiles of *GZMH* and *GZMK*.

**Figure 3:**
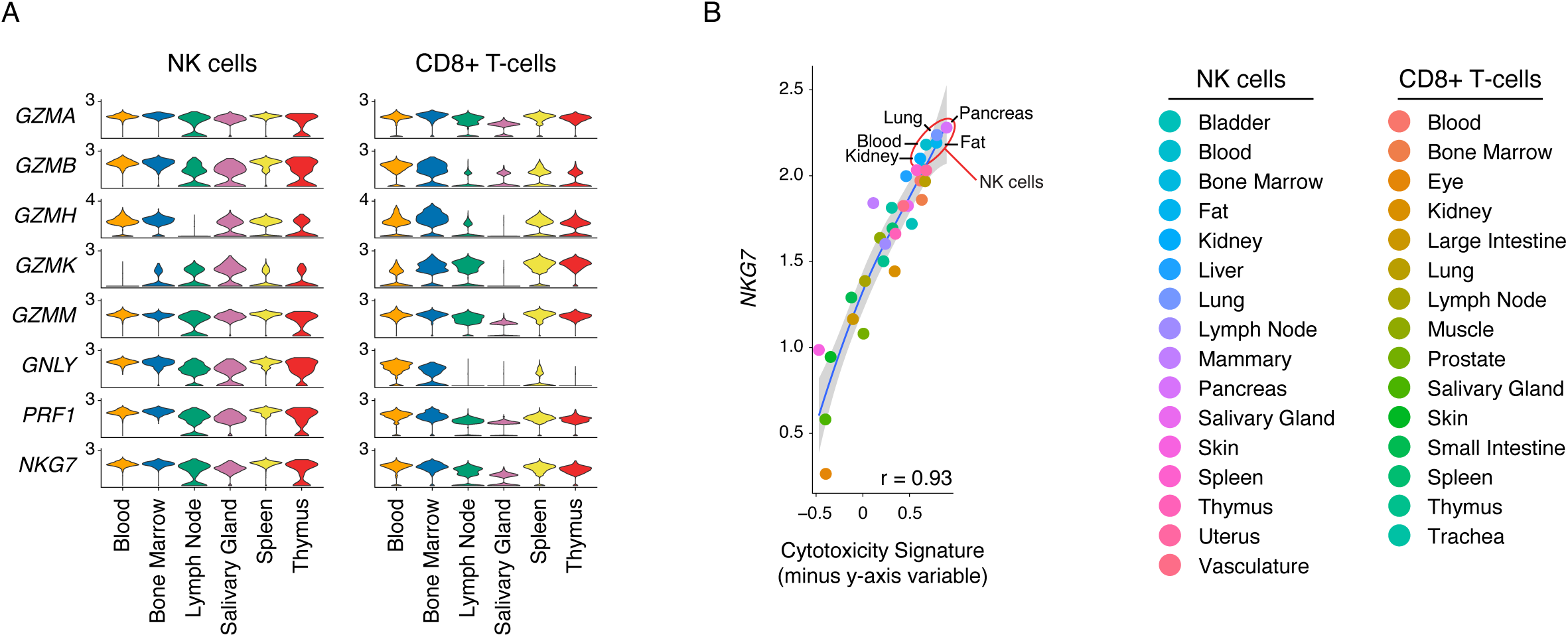
NKG7 correlates with cytotoxicity across organs. (A) Violin plots showing the gene expression of cytotoxic molecules in NK (left) and CD8+ T-cells (Right) across major immunological organs. (B) Scatterplots of the correlation between NKG7 gene expression and cytotoxicity score within NK or CD8+ T-cells across all organs of the tabula sapiens dataset. Pearson correlation is displayed and the shaded area represents the 95% CI.

Similarly, in CD8+ T-cells, *NKG7* was consistently detected, although there was a notable drop in expression observed in salivary gland CD8+ T-cells. Across both NK and CD8+ T-cells, *GZMA*, *GZMM*, *PRF1*, and *NKG7* were consistently expressed and detected regardless of tissue. Importantly, *GZMB* had variable expression within CD8+ T-cells and was minimally detected in lymph node, salivary gland, and thymic tissues. As such, this robust expression of *NKG7* extended across all tissues of the Tablua Spaiens dataset and *NKG7* gene expression was found to be strongly correlated (r = 0.93) with the cytotoxicity signature within NK and CD8+ T-cells from various tissues (Figure 3B). In contrast, *GZMB* showed a poorer correlation with cytotoxicity across tissues with a pearson r = 0.72 (data not shown). Therefore, these data demonstrate that NKG7 robustly captures cytotoxicity within traditionally cytotoxic populations regardless of tissue of origin.

### NKG7 is consistently expressed in polyfunctional cytotoxic cells

The expression of cytotoxic molecules is rarely analyzed at single-cell resolution, making it difficult to distinguish between polyfunctional and monofunctional cytotoxic cells. To address this, we utilized single-cell data to examine the co-expression patterns of cytotoxic molecules. Transcriptionally, it was observed that a considerable proportion (∼27%) of pro-inflammatory NK-Bright cells co-expressed *GZMA*, *GZMK*, *GNLY*, *PRF1*, and *NKG7* (Figure 4A). In contrast, the cytotoxic NK-Dim subset consistently co-expressed *GZMA*, *GZMB*, *GNLY*, *PRF1*, and *NKG7*, with polyfunctional populations of NK-Dim cells differing in their co-expression of *GZMH* and *GZMM*. Interestingly, circa 20% of dnT cells were observed to exhibit solitary expression of *GZMK,* while a small subset of dnT cells co-expressed *GZMK* with either *GZMA*, *GZMM*, or both. An appreciable frequency of CD8-Naïve and CD8-CM populations subsets are characterized by the singular expression of cytotoxic molecules, even though these subsets are largely not cytotoxic. In contrast, cytotoxic subsets such as CD8-EM and CD4-CTLs demonstrated diverse co-expression patterns. For example, while the top five most abundant co-expression patterns for CD8-EM and CD4-CTLs consistently contained *GZMA* and *NKG7*, there was variable usage of other cytotoxic molecules. Within CD8-EM cells, all remaining cytotoxic molecules were variably expressed. However, within CD4-CTLs, there was consistent expression of *GZMH* and *GNLY*. While *GZMK* was absent from the top five most abundant CD4-CTL cell patterns. Notably, three of the top five most abundant CD8-EM and CD4-CTL patterns did not contain *PRF1*. These findings highlight that *NKG7* is consistently observed within polyfunctional cytotoxic subsets across diverse immune cell populations.

**Figure 4:**
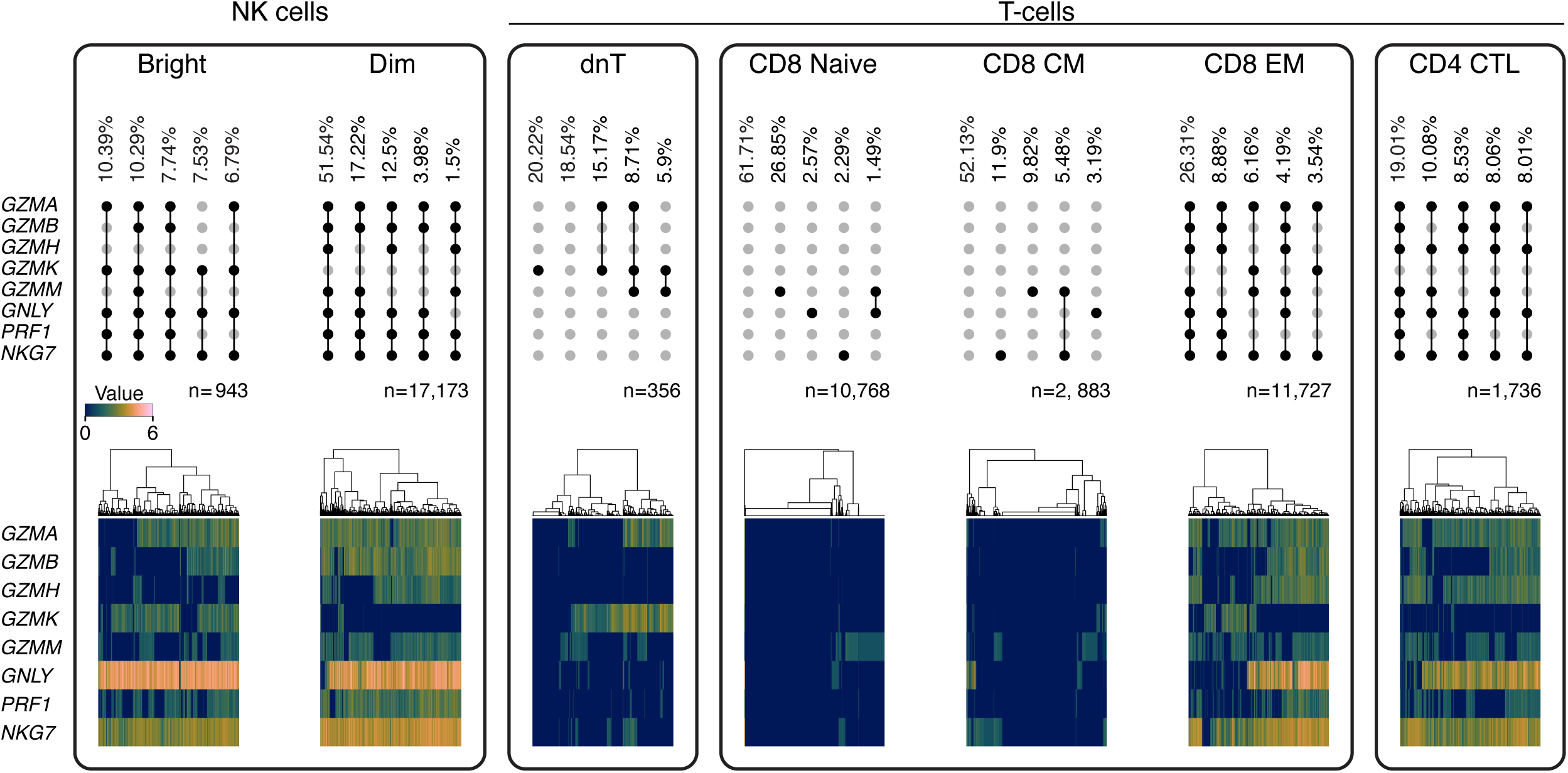
NKG7 is consistently co-expressed with other cytotoxic molecules. Co-expression pattern of cytotoxic molecules across NK and T-cell subsets. Upset plots (top row) demonstrate the co-expression pattern and frequency observed. Meanwhile, heatmaps (bottom row) demonstrate the expression profile of individual cells. n value represents the number of cells of a particular subset included in the analysis.

### NKG7 identifies cytotoxic tumor-infiltrating cells

NKG7 is consistently expressed across tissues and as a core constitute of the polyfunctional cytotoxic program. However, it is known that the tumor microenvironment (TME) can drastically alter expression patterns of infiltrating cytotoxic immune cells. Therefore, we next sought to evaluate cytotoxic molecule expression dynamics in tumor infiltrating immune cells. To investigate this, we utilized the Tumor Immune Cell Atlas (Nieto et al., 2021). This atlas contains tumor-infiltrating cells (n = 314,679) from 177 patients spanning 12 tumor subtypes (Supplementary Figure 2A). Scoring cells within the dataset for overall cytotoxicity reveals two areas of dense signal, corresponding to the CD8 T-cell and NK cell populations (Figure 5A–C). Interestingly, cells annotated as terminally exhausted CD8 T-cells show high levels of expression of cytotoxicity genes. Consistent with previous observations, *NKG7* gene expression strongly correlates with overall cytotoxicity (Figure 5D). Indeed, *NKG7* is the strongest correlate of cytotoxicity across tumor-infiltrating immune cells. Surprisingly, *GZMH* showed the second highest correlation with cytotoxicity while *GZMB* scored poorly. The low correlation of *GZMB* expression with overall cytotoxicity signature is driven by the unique and singular expression of *GZMB* in pDC cells. However, even in the absence of pDC cells, *GZMB* expression was a poorer correlate of cytotoxicity score than *NKG7* gene expression (Supplementary Figure 2B). NK cells are a potent cytotoxic population but are poorly captured within the Tumor Immune Cell Atlas (*n* = 9,496). Therefore, we utilized a pancancer NK cell atlas (*n* = 34,900) to further investigate cytotoxic molecule expression within tumor-infiltrating NK cells (Tang et al., 2023). The NK cell subsets referenced in this study were pre-defined within the pancancer NK cell atlas dataset. Within these tumor-infiltrating NK subsets, NKG7 was consistently expressed (Figure 5E). Additionally, *NKG7* was highly expressed in tumor-infiltrating NK cells from all disease subsets analysed (Supplementary Figure 2C). As such, *NKG7* gene expression positively correlated with overall cytotoxicity in tumor-infiltrating NK cells (Figure 5F). *NKG7* was a stronger correlate for cytotoxicity than all other markers except *GZMA* and *PRF1* (Supplementary Figure 2D). Taken together, these data demonstrate that *NKG7* expression is maintained on tumor-infiltrating immune cells in numerous malignancies. Furthermore, *NKG7* positively correlates with overall cytotoxicity, outperforming traditional markers of cytotoxicity such as *GZMB*.

**Figure 5:**
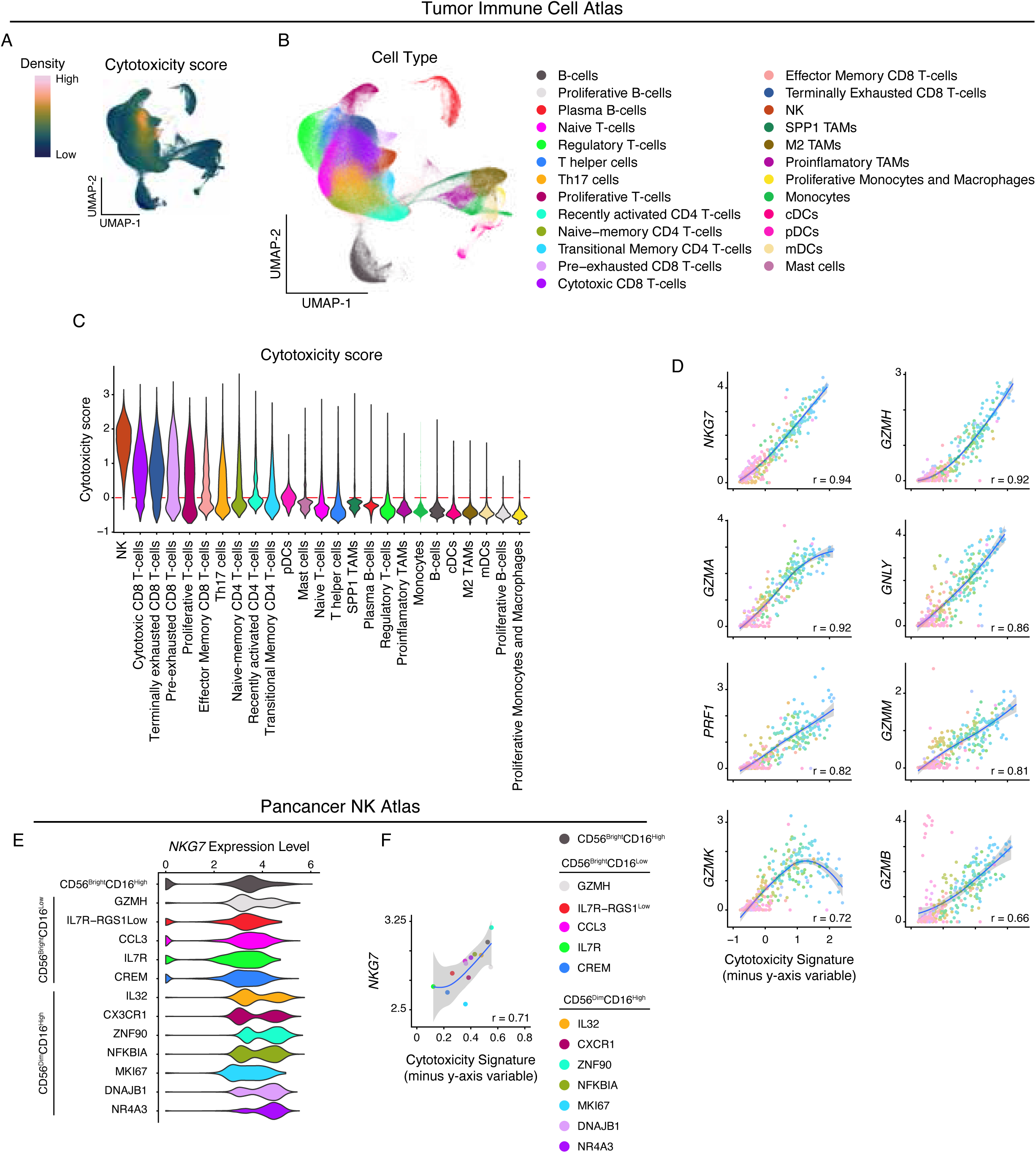
NKG7 correlates with cytotoxicity in tumor-infiltrating immune subsets. (A) Density of cytotoxicity score expression overlaid on UMAP coordinates of the tumor immune cell atlas. (B) UMAP plot of the different cell subsets identified within the tumor immune cell atlas. (C) Violin plots showing the cytotoxicity score of various cell subsets within tumor immune cell atlas. (D) Scatterplot of the correlation between y-axis gene expression and cytotoxicity signature. Pearson correlation is displayed and the shaded area represents the 95% CI. (E) UMAP plot highlighting the NK subsets identified within the pancancer NK atlas (left). Violin plot depicts the expression of NKG7 across various NK subsets within the pancancer NK atlas. (F) Scatterplot of the correlation between NKG7 or Granzyme B (GZMB) expression and cytotoxicity score across NK subsets. Pearson correlation is displayed and the shaded area represents the 95% CI.

## Discussion

NKG7 is an emerging component of the cytotoxic machinery, yet its functional roles and expression dynamics remain poorly understood. To address this gap, we conducted a comprehensive analysis of NKG7 expression alongside other core components of cytotoxic granules, providing insights into its reliability as a cytotoxicity marker across diverse contexts. As such, it was revealed that NKG7 is a robust and reliable marker of cytotoxicity across healthy and disease contexts. NKG7 is consistently expressed across various cytotoxic immune subsets and immunologically relevant tissues. Notably, *NKG7* demonstrated superior reliability compared to traditional markers such as granzyme B (GZMB) and perforin, particularly in distinguishing functional cytotoxicity. For example, while plasmacytoid dendritic cells (pDCs) express solitary GZMB, this alone does not reflect engagement with the broader cytotoxic program. In contrast, NKG7 expression correlates consistently with active cytotoxic machinery across immune subsets. Cytotoxic tumor-infiltrating lymphocytes (TILs) are critical determinants of patient prognosis, with higher frequencies consistently correlating with improved survival across malignancies (Brummel et al., 2023). Transcript-based signatures, such as the cytolytic activity score (GZMA+ PRF1+), have been developed to quantify TIL cytotoxicity and predict disease outcomes (Narayanan et al., 2018; Rooney et al., 2015; Wakiyama et al., 2018). Notably, NKG7 outperforms many traditional markers, offering improved resolution in capturing functional cytotoxicity. In line with this, *NKG7* expression has been reported to be associated with favorable clinical outcomes and is important for effective T-cell immunity (Lelliott et al., 2022b; Li et al., 2022; Ng et al., 2020; Wen et al., 2022). This growing recognition underscores the importance of understanding NKG7’s integration into broader cytotoxicity programs.

Our analysis revealed the underappreciated heterogeneity of cytotoxic molecule usage. For instance, pro-inflammatory (NK-Bright) and cytotoxic (NK-Dim) cells exhibit distinct usage of *GZMK* and *GZMH*, reflecting functional specialization within cytotoxic programs (Jiang et al., 2011). Similarly, tissue-specific cytotoxic programs in NK and CD8+ T-cells demonstrated substantial variability in granzyme expression, which likely reflects differences in subset composition and tissue reprogramming. For example, differences in granzyme expression among NK subsets may reflect tissue-resident programming, while maturation state influences granzyme patterns in CD8+ T-cells. Despite these variations, NKG7 expression remained consistent across tissues and subsets, further supporting its role as a universal cytotoxicity marker.

Single-cell analysis revealed cytotoxic cells lacking *PRF1,* suggesting alternative pathways for granzyme activity. Mechanisms mediating granzyme entry, such as mannose-6-phosphate receptor or serglycin-mediated transport, may underlie these observations (Raja et al., 2005; Veugelers et al., 2006). Notably, NKG7 expression was preserved in these populations, suggesting its potential role in cytotoxic programs that operate independently of PRF1. However, the extent to which these pathways operate independently of PRF remains unclear. Experimental evidence indicates that perforin (and not granzymes) is essential for effective tumor control (Smyth et al., 2003). This suggests that perforin-negative populations may be leveraging granzymes for non-cytotoxic roles, such as extracellular matrix (ECM) remodeling. Granzymes contribute to ECM remodeling by cleaving proteins such as fibronectin and laminin, with complex effects on tumor progression. While ECM degradation can enhance immune infiltrating and induce anoikis, it may also promote metastasis by weakening cell adhesion. Additionally, ECM breakdown releases immunomodulatory cytokines and chemotactic fragments (Boivin et al., 2009). Demonstrating the dual roles of granzymes in immune regulation and cancer progression. Despite this complexity, NKG7 consistently marks cytotoxic cells, irrespective of whether engaged in cytotoxic or non-cytotoxic functions.

NKG7 consistently emerged as a central component of the cytotoxic molecule expression program across NK subsets and effector states such as CD8-EM and CD4-CTL, identifying cytotoxic cells regardless of their molecular programs. NKG7 also functioned to demarcate CD4-CTLs, a subset with emerging relevance in anti-tumor responses. Indeed, NKG7 is frequently observed in gene signatures of CD4-CTLs across numerous disease contexts (Cachot et al., 2021; Maehara et al., 2020; Oh et al., 2020; Preglej & Ellmeier, 2022; Zhang et al., 2020; Zheng et al., 2017). Additionally, NKG7 was consistently highly correlated with cytotoxicity within TILs, surpassing other canonical markers of cytotoxicity. This underscores its stability and reliability as an effective correlate of cytotoxicity even in the immunosuppressive tumor microenvironment. As such, NKG7 represents a central and versatile marker of cytotoxicity, distinguished by its consistency across immune subsets, tissues, and disease contexts. Its stability and integration into diverse cytotoxic programs position NKG7 as a valuable tool for both research and clinical applications.

## Methods

### Cell culture and transfection of HEK293T cells

Human embryonic kidney 293 cells (HEK293T) were cultured in DMEM medium, supplemented with GLUTAMax (Gibco, # 61965-026), 10% (vol/vol) Fetal calf serum (FCS) (Gibco, # A5256801) and 100 units/mL of penicillin G and 100 ug/mL of streptomycin sulfate (Gibco, # 15140-122) at 37°C, 5% CO_2_.

For the transfection, HEK293 cells were seeded in 6-well plate (1×10^6^ cells/well) and incubated overnight. On the next day, cells were transfected via calcium phosphate method, as previously described (“Calcium Phosphate–Mediated Transfection of Eukaryotic Cells,” 2005), so that they could overexpress mNeon-NKG7, or 3xFLAG-TIA-1. Plasmids were purchased from BioCat and sequences can be found as following: mNeonGreen-Linker-NKG7(165aa), cloning vector pcDNA3.0, cloning sites NotI(GCGGCCGC)-XhoI(CTCGAG); sequence: GCGGCCGCCACCATGGTGAGCAAGGGCGAGGAGGATAACGCCTCTCTCCCAGCGACACATGAGTTACACATCTTTGGCTCCATCAACGGTGTGGACTTTGACATGGTGGGTCAGGGCACCGGCAATCCAAATGATGGTTATGAGGAGTTAAACCTGAAGTCCACCAAGGGTGACCTCCAGTTCTCCCCCTGGATTCTGGTCCCTCATATCGGGTATGGCTTCCATCAGTACCTGCCCTACCCTGACGGGATGTCGCCTTTCCAGGCCGCCATGGTAGATGGCTCCGGATACCAAGTCCATCGCACAATGCAGTTTGAAGATGGTGCCTCCCTTACTGTTAACTACCGCTACACCTACGAGGGAAGCCACATCAAAGGAGAGGCCCAGGTGAAGGGGACTGGTTTCCCTGCTGACGGTCCTGTGATGACCAACTCGCTGACCGCTGCGGACTGGTGCAGGTCGAAGAAGACTTACCCCAACGACAAAACCATCATCAGTACCTTTAAGTGGAGTTACACCACTGGAAATGGCAAGCGCTACCGGAGCACTGCGCGGACCACCTACACCTTTGCCAAGCCAATGGCGGCTAACTATCTGAAGAACCAGCCGATGTACGTGTTCCGTAAGACGGAGCTCAAGCACTCCAAGACCGAGCTCAACTTCAAGGAGTGGCAAAAGGCCTTTACCGATGTGATGGGCATGGACGAGCTGTACAAGGGGTCTGGTGGCAGTGGAGGGGGATCCATGGAGCTCTGCCGGTCCCTGGCCCTGCTGGGGGGCTCCCTGGGCCTGATGTTCTGCCTGATTGCTTTGAGCACCGATTTCTGGTTTGAGGCTGTGGGTCCCACCCACTCAGCTCACTCGGGCCTCTGGCCAACAGGGCATGGGGACATCATATCAGGCTACATCCACGTGACGCAGACCTTCAGCATTATGGCTGTTCTGTGGGCCCTGGTGTCCGTGAGCTTCCTGGTCCTGTCCTGCTTCCCCTCACTGTTCCCCCCAGGCCACGGCCCGCTTGTCTCAACCACCGCAGCCTTTGCTGCAGCCATCTCCATGGTGGTGGCCATGGCGGTGTACACCAGCGAGCGGTGGGACCAGCCTCCACACCCCCAGATCCAGACCTTCTTCTCCTGGTCCTTCTACCTGGGCTGGGTCTCAGCTATCCTCTTGCTCTGTACAGGTGCCCTGAGCCTGGGTGCTCACTGTGGCGGTCCCCGTCCTGGCTATGAAACCTTGTGACTCGAG 3xFLAG-Linker-TIA1(386aa), cloning vectori: pcDNA3.0, cloning sites NotI(GCGGCCGC)-XhoI(CTCGAG); sequence: GCGGCCGCCACCATGGACTATAAGGACCACGACGGAGACTACAAGGATCATGATATTGATTACAAAGACGATGACGATAAGGGGTCTGGTGGCAGTGGAGGGGGATCCATGGAGGACGAGATGCCCAAGACTCTATACGTCGGTAACCTTTCCAGAGATGTGACAGAAGCTCTAATTCTGCAACTCTTTAGCCAGATTGGACCTTGTAAAAACTGCAAAATGATT ATGGATACAGCTGGAAATGATCCCTATTGTTTTGTGGAGTTTCATGAGCATCGTCATGCAGCTGCAGCATTAGCTGCTATGAATGGACGGAAGATAATGGGTAAGGAAGTCAAAGTGAATTGGGCAACAACCCCTAGCAGTCAAAAGAAAGATACAAGCAGTAGTACCGTTGTCAGCACACAGCGTTCACAAGATCATTTCCATGTCTTTGTTGGTGATCTCAGCCCAGAAATTACAACTGAAGATATAAAAGCTGCTTTTGCACCATTTGGAAGAATATCAGATGCCCGAGTGGTAAAAGACATGGCAACAGGAAAGTCTAAGGGATATGGCTTTGTCTCCTTTTTCAACAAATGGGATGCTGAAAACGCCATTCAACAGATGGGTGGCCAGTGGCTTGGTGGAAGACAAATCAGAACTAACTGGGCAACCCGAAAGCCTCCCGCTCCAAAGAGTACATATGAGTCAAATACCAAACAGCTATCATATGATGAGGTTGTAAATCAGTCTAGTCCAAGCAACTGTACTGTATACTGTGGAGGTGTTACTTCTGGGCTAACAGAACAACTAATGCGTCAGACTTTTTCACCATTTGGACAAATAATGGAAATTCGAGTCTTTCCAGATAAAGGATATTCATTTGTTCGGTTCAATTCCCATGAAAGTGCAGCACATGCAATTGTTTCTGTTAATGGTACTACCATTGAAGGTCATGTTGTGAAATGCTATTGGGGCAAAGAAACTCTTGATATGATAAATCCCGTGCAACAGCAGAATCAAATTGGATATCCCCAACCTTATGGCCAGTGGGGCCAGTGGTATGGAAATGCACAACAAATTGGCCAGTATATGCCTAATGGTTGGCAAGTTCCTGCATATGGAATGTATGGCCAGGCATGGAACCAGCAAGGATTTAATCAGACACAGTCTTCTGCACCATGGATGGGACCAAATTATGGAGTGCAACCGCCTCAAGGGCAAAATGGCAGCATGTTGCCCAATCAGCCTTCTGGGTATCGAGTGGCAGGGTATGAAACCCAGTGACTCGAG. Cells were collected after 24 hours from the transfection, washed twice in PBS and then lysed to extract proteins, as described in the next paragraph.

### Western blots

Cells were lysed using RIPA lysis buffer supplemented with protease and phosphatase inhibitors diluted 1:100 (cat. 5872, Cell Signaling,). The total protein concentration was determined with the Protein Assay Dye Reagent (cat. 5000006, Bio-Rad,). For each sample, 20 µg of protein were prepared in Laemmli buffer and separated on 15 % polyacrylamide gels via SDS-PAGE. After electrophoresis, proteins were transferred to nitrocellulose blotting membranes (cat. 10600004, Cytiva) in a wet blotting Mini Trans-Blot® Cell system (Bio-Rad). The membranes were incubated with 5 % (w/v) bovine serum albumin (BSA) (cat. 8076, Carl Roth) in TRIS-buffered saline with 0.05 % (v/v) of Tween 20 (TBS-T) for 1 hour at room temperature followed by incubation with primary antibodies overnight at 4 °C. All primary antibodies were diluted in 5 % (w/v) BSA in TBS-T with following dilutions: anti-NKG7 (Beckman Coulter, #IM2550, 2G9A10F5, 1:200), anti-TIA-1 (clone EPR9304, 1:1000, cat. ab140595, abcam), anti-FLAG (clone M2, 1:500, cat. F3165, Sigma-Aldrich), anti-β-actin (clone 13E5, 1:1000, cat. 4970, Cell Signaling Technology) anti-mNeonGreen (clone 32F6, 1:200, cat. 32f6, Proteintech), anti-NKG7 (polyclonal, 1:1000, cat. 65507, Cell Signaling Technology), anti-FLAG (clone L5, 1:500, cat. NBP1-06712, Novus Biologicals), anti-mNeonGreen (polyclonal, 1:1000, cat. 53061, Cell Signaling Technology). After overnight incubation with primary antibodies, the membranes were washed three times with TBS-T for 5 minutes each and incubated with secondary antibodies for 1 hour at room temperature. All secondary antibodies were used 1:15000 diluted in 5 % (w/v) BSA in TBS-T: IRDye® 800CW Goat anti-Mouse (cat. 926-32210, LI-COR), IRDye® 800CW Donkey anti-Rabbit (cat. 926-32213, LI-COR), IRDye® 680RD Goat anti-Rat (cat. 926-68076, LI-COR), IRDye® 680RD Goat anti-Rabbit (cat. 926-68071, LI-COR), IRDye® 680RD Donkey anti-Mouse (cat. 926-68072, LI-COR). After incubation with secondary antibodies, the membranes were washed three times with TBS-T for 5 minutes each and the protein bands were imaged using the Odyssey® CLx system (LI-COR).

### Peripheral Blood Mononuclear Cell (PBMC) dataset

#### Dataset and Preprocessing

The PBMC dataset published and described in (Hao et al., 2021) was downloaded from https://atlas.fredhutch.org/nygc/multimodal-pbmc/. This dataset contains PBMCs from eight healthy donors collected at three timepoints relative to HIV vaccination: pre-vaccination, 3 days post-vaccination, and 7 days post-vaccination. In total, it includes 161,764 cells and 20,957 features, comprising RNA expression and 228 CITE-seq antibody-derived tags (ADTs).

The original study performed quality control, cell type annotation, and multimodal integration, using both RNA and ADT data for cell type identification and dimensionality reduction (see Hao et al., 2021 for details). For our analysis, we focused only on cells from the pre-vaccination timepoint. Cells annotated as doublets at the celltype.l1 annotation level were removed, reducing the dataset to 161,159 cells. All downstream analysis utilized the dimensionlity reduction and cell type classifications provided by the original dataset authors.

#### Custom Cell Type Annotation

The dataset authors provided cell type annotations at multiple levels of resolution, with celltype.l1 represeting braod cell classificatons and celltype.l3 offering more detailed subtype annotations (Hao et al., 2021). To simplify and standardize labeling, we re-annotated the dataset by consolidating celltype.l3 categories into broader, more interpretable groups. This post-processing step retained the structure of the original dataset but grouped similar populations under unified labels. Specifically, the following groupings were applied. All B-cell subtypes, including “B intermediate lambda,” “B naive kappa,” “B intermediate kappa,” “B memory kappa,” “B naive lambda,” and “B memory lambda,” were combined into a single category labeled “B_cells.” The “Plasma” and “Plasmablast” subtypes were grouped together as “Plasmablasts.” The NK subsets labeled “NK_1” through “NK_4” were merged into a single category called “NK_Dim,” while “Treg Naive” and “Treg Memory” were combined under the label “Treg.” The “dnT_1” and “dnT_2” subsets were consolidated into the “dnT” category. Similarly, the dendritic cell subsets “cDC2_1,” “cDC2_2,” and “ASDC_mDC” were grouped as “cDC2,” while “ASDC_pDC” and “pDC” were merged into the “pDC” category. The gamma-delta T-cell subsets “gdT_2,” “gdT_3,” and “gdT_4” were combined into a single “gdT” category, whereas “gdT_1” was retained as “gdT_V9D2.“

For CD4+ T-cell subsets, “CD4 TCM_1” through “CD4 TCM_3” were grouped as “CD4_CM,” and “CD4 TEM_1” through “CD4 TEM_4” were combined as “CD4_EM.” For CD8+ T cells, “CD8 Naive” and “CD8 Naive_2” were merged into “CD8_Naive,” while “CD8 TEM_1” through “CD8 TEM_6” were consolidated under “CD8_EM.” Additionally, “CD8 TCM_1” through “CD8 TCM_3” were grouped as “CD8_CM.” To maintain consistency, formatting adjustments were applied, including replacing spaces with underscores and standardizing nomenclature for “Platelets” and “Prolif” (proliferating cells). This post-processing step resulted in 28 distinct cell populations used for downstream analyses.

### Tabula Sapiens dataset

#### Dataset and Preprocessing

The TS_immune dataset from the Tabula Sapiens project was downloaded from figshare (https://figshare.com/projects/Tabula_Sapiens/100973). This dataset is a comprehensive single-cell RNA sequencing atlas of 58,870 genes across 264,824 cells from 24 different anatomical sites (THE TABULA SAPIENS CONSORTIUM, 2022). These data were derived from both 10x Genomics and Smart-seq2 technologies, with the majority of cells originating from 10x Genomics. For this analysis, we filtered the data to retain only cells processed with 10x Genomics, resulting in 249,961 cells. Donors with a low cell count (fewer than 50 cells) were excluded, specifically TSP3 (2 cells), TSP12 (48 cells), and TSP13 (0 cells), leaving 12 donors in the final dataset.

#### Data Annotation and Cleaning

The cell type annotations provided with the dataset were cleaned and formatted to resolve inconsistencies and correct misspellings. Details of this cleaning process are available in the GitHub repository associated with this manuscript. See code availability for more information.

#### Normalization and Integration

The dataset was split into individual layers by donor, and data normalization was then performed on each donor separately. Data normalization was performed using the **SCTransform** function (vst.flavor = “v2”, method = glmGamPoi). Principal component analysis (PCA) was conducted using **RunPCA** with default parameters. The datasets were then integrated using the **IntegrateLayers** function with canonical correlation analysis (CCA) as the integration method. This step utilized PCA dimensions from the SCTransform-normalized data.

#### Dimensionality Reduction and Imputation

Uniform manifold approximation and projection (UMAP) was calculated using **RunUMAP** on 30 dimensions of the integrated CCA reduction. For specific visualizations, imputed values were used to enhance interpretability. Imputation was performed using **RunALRA,** which increased the proportion of nonzero entries in the data matrix from 16.19% to 49.97%.

### Tumor Immune Cell Atlas (TICAtlas)

The TICAtlas dataset was downloaded from Zenodo (https://zenodo.org/records/4263972) and is described in detail in the accompanying publication (Nieto et al., 2021). The dataset originally contained 92,256 features across 317,111 cells, representing 13 tumor types from 181 patients and annotated with 25 cell types. For the analysis, cells from the ovarian cancer (OC) subtype were removed due to the low number of cells available (2,432 total). Consequently, the analyzed dataset included data from 177 patients across 12 tumor subtypes.

### Pancancer NK atlas

The NK Atlas dataset was downloaded from Zenodo (https://zenodo.org/records/8275845) and contained expression data for 13,493 genes across 142,304 cells and is described in detail in the accompanying publication (Tang et al., 2023). Data was filtered to include only tumor-derived cells, resulting in a final dataset of 34,900 cells across 24 tumor types and 13 NK subtypes as defined in the original dataset.

### Software and Versions

All analyses were conducted using the R programming environment (v4.4.0) (R Core Team, 2023) on a platform of x86_64-apple-darwin20 running macOS Ventura 13.0. The primary tools included the **Seurat** package (v5.1.0) (Hao et al., 2023) for data normalization, integration, dimensionality reduction, and visualization, with additional functionality provided by **SeuratDisk** (v0.0.0.902) (Hoffman, Paul, 2023), **SeuratWrappers** (v0.3.2) (Satija, Butler, et al., 2023), and **SeuratObject** (v5.0.2) (Satija, Hoffman, et al., 2023).

Visualization and data processing were performed using **ggplot2** (v3.4.4) (Wickham, 2016), **dplyr** (v1.1.4) (Wickham et al., 2023), **scCustomize** (v2.1.2) (Marsh, 2023), and **Nebulosa** (v1.14.0) (Alquicira-Hernandez & Powell, 2021). Heatmaps and density plots utilized the “batlow” color scheme, accessed through the **scico** package (v1.5.0) (Pedersen & Crameri, 2023).

### Correlation Analysis

To perform the correlation analysis, expression data was first aggregated using the **AggregateExpression** function. Module score was calculated using **AddModuleScore** function across a number of gene sets. The gene sets were defined by iteratively removing one marker at a time from the full list of cytotoxicity markers. Specifically, the full cytotoxicity signature consisted of the following markers: GZMA, GZMB, GZMH, GZMK, GZMM, GNLY, PRF1, and NKG7. Pairwise correlation plots were generated using the **FeatureScatter** function to compare the module scores from each gene set with the corresponding marker that was excluded from the set.

### Bulk RNAseq dataset

The “Monaco” dataset (Monaco et al., 2019) of RNA transcript abundance across human peripheral blood immune cells was downloaded from the Human Protein Atlas (https://www.proteinatlas.org). Data was plotted using GraphPad Prism (v10.0.3, GraphPad Software, Boston, Massachusetts USA, www.graphpad.com).

### PBMCs isolation

Peripheral blood from healthy donors was provided by the Institute for Experimental Hematology and Transfusion Medicine at the University Hospital Bonn, Bonn, Germany. PBMCs were isolated from peripheral blood by Ficoll-Paque (Cytiva, cat. 17144003) density gradient, following the standard protocol. NK cells were isolated from PBMCs using the EasySep Human NK cell Isolation Kit (Stemcell Technologies, Cat. 17955). After being isolated, PBMCs and NK cells were processed for flow cytometry staining.

### Flow cytometry

Flow cytometry staining was performed in 96-well round bottom microplates (cat 92697, TPP), at room temperature and protected from light. Cells were firstly washed twice in PBS, then incubated with 50 uL TruStain FcX (1:200, cat. 422302, Biologend) and Live/Dead Blue Fixable dye (1:1000, cat. L23105, Invitrogen), for 15 minutes at room temperature. After washing twice in PBS cells were then incubated with 50 uL of a cocktail of fluorescence-conjugated antibodies recognizing surface molecules, for 20 min, at room temperature. The cocktail for the surface staining contained the following antibodies at the indicated dilution: BUV395 CD8 (clone RPA-T8, 1:200, cat. 563796, BD Biosciences), BUV496-CD16 (clone 3G8, 1:200, cat. 612944, BD Biosciences), BUV563-CD56 (clone NCAM16, 1:200, cat. 612928, BD Biosciences), BV570-CD45RA, (clone HI100, 1:100, cat. 304130, BioLegend), BV650-CD4 (clone OKT4, 1:200, cat. 317435, BioLegend), BV785-CD62L (clone DREG-56, 1:200, cat. 304829, BioLegend), APC-Cy7-CD3 (clone SK7, 1:100, cat. 557832, BD Biosciences). After the incubation with the antibody cocktail, samples were washed twice in PBS and then incubated with 100 uL of eBioscience Foxp3/Transcription Factor Staining Buffer Set (ThermoFisher, cat. 00-5523-00), for 15 minutes. Cells were then washed twice with wash buffer and incubated for 30 minutes with 50 uL of a cocktail of fluorescence-conjugated antibodies reactive against intracellular targets. The cocktail contained the following antibodies: Pacific Blue-Granzyme A (clone CB9, 1:50, cat. 507207, BioLegend), BV711-Perforin (clone dG9, 1:50, cat. 308130, BioLegend), AF488-Granzyme H (clone E3H7W, cat. 23455S, Cell Signaling Technologies), PE-NKG7 (clone 2G9A10F5, 1:100, cat. IM3293, Beckman Coulter), PE-Dazzle-594-Granzyme B (clone QA18A28, 1:50 cat. 396427, BioLegend), PE-Cy7-Granzyme K (clone GM26E7, 1:50, cat. 370515, BioLegend), AF-647-Granzyme M (clone 4B2G4, 1:50, cat. 566996, BD Biosciences), AF700-Granulysin (clone B-L38, 1:50, cat. NBP3-18104AF700, Novus Bio). As controls, a second antibody cocktail containing the following IgG controls: BV711-Mouse IgG2k (clone MPC-11, cat. 400354, BioLegend), AF88-Rabbit IgG (polyclonal, cat. 4340S, Cell Signaling Technologies), PE-Dazzle-594-Rat IgG1k (clone RTK2071, cat. 400445, BioLegend), PE-Cy.7-Mouse IgG1k (clone MOPC-21, cat. 400125, BioLegend), AF647-Mouse IgG1k (clone P3.6.2.8.1, cat. 51-4714-81, eBioscience), AF700-Mouse IgG1k (clone MOPC-21, cat. 400143, BioLegend). After the incubation, all the samples were washed in FACS Buffer (PBS, 0.02% (vol/vol) FCS, 5 mM EDTA and 0.01%), and then fixed in 4% PFA (HistoFix, Roth, cat. P087.5). Cells were washed twice in FACS buffer and stored at +4°C, until acquisition. Samples were acquired on a Sony ID7000 7 lasers. Data were analyzed using FlowJo™ Software (v10.9.0, BD Life Sciences).

### Statistical analysis

Statistical analyses were performed using GraphPad Prism (v10.0.3, GraphPad Software) or R programming environment (v4.4.0) (R Core Team, 2023). The specific statistical test used for each analysis are indicated in the corresponding figure legends.

## Figure preparation

Figures were arranged and formatted using Adobe Illustrator (v27.5, Adobe Inc.) and/or GraphPad Prism (v10.0.3, GraphPad Software)

## Materials availability

This study did not generate any new materials

## Data and code availability

The datasets analysed in this study were obtained from https://www.proteinatlas.org (Bulk RNAseq dataset), https://atlas.fredhutch.org/nygc/multimodal-pbmc/ (PBMC dataset), https://figshare.com/projects/Tabula_Sapiens/100973 (Tablua Sapiens dataset), https://zenodo.org/records/4263972 (Tumor Immune Cell Atlas dataset), and https://zenodo.org/records/8275845 (Pancancer NK Atlas dataset). All code used to generate figures can be found under the relevant repository at https://github.com/BaldLab. All other data is available upon request.

## Acknowledgements

The authors thank the Flow Cytometry Core Facility (FCCF) of the Medical Faculty at the University of Bonn for providing help, services, and devices.

## Authors contributions

**Roberta Turiello:** Conceptualization, Validation, Formal analysis, Investigation, Data curation, Visualization, Writing - Original Draft, Writing - Review & Editing

**Susanna S. Ng:** Conceptualization, Validation, Formal analysis, Investigation, Data curation, Visualization, Writing - Original Draft, Writing - Review & Editing

**Elisabeth Tan**: Investigation, Data Curation, Writing - Review & Editing

**Gemma van der Voort**: Investigation, Data Curation, Writing - Review & Editing

**Nazhifah Salim**: Investigation, Data Curation, Writing - Review & Editing

**Michelle Yong**: Investigation, Writing - Review & Editing

**Malika Khassenova**: Methodology, Investigation

**Johannes Oldenburg:** Resources

**Heiko Rühl**: Resources

**Jan Hasenaur**: Writing - Review & Editing

**Laura Surace**: Writing - Review & Editing

**Marieta Toma**: Resources

**Tobias Bald:** Writing - Review & Editing

**Michael Hölzel:** Conceptualization, Supervision, Project administration, Funding acquisition, Writing - Review & Editing

**Dillon Corvino:** Conceptualization, Software, Formal analysis, Data curation, Visualization, Supervision, Project administration, Writing - Original Draft, Writing - Review & Editing

---

**Supplementary Figure 1.**
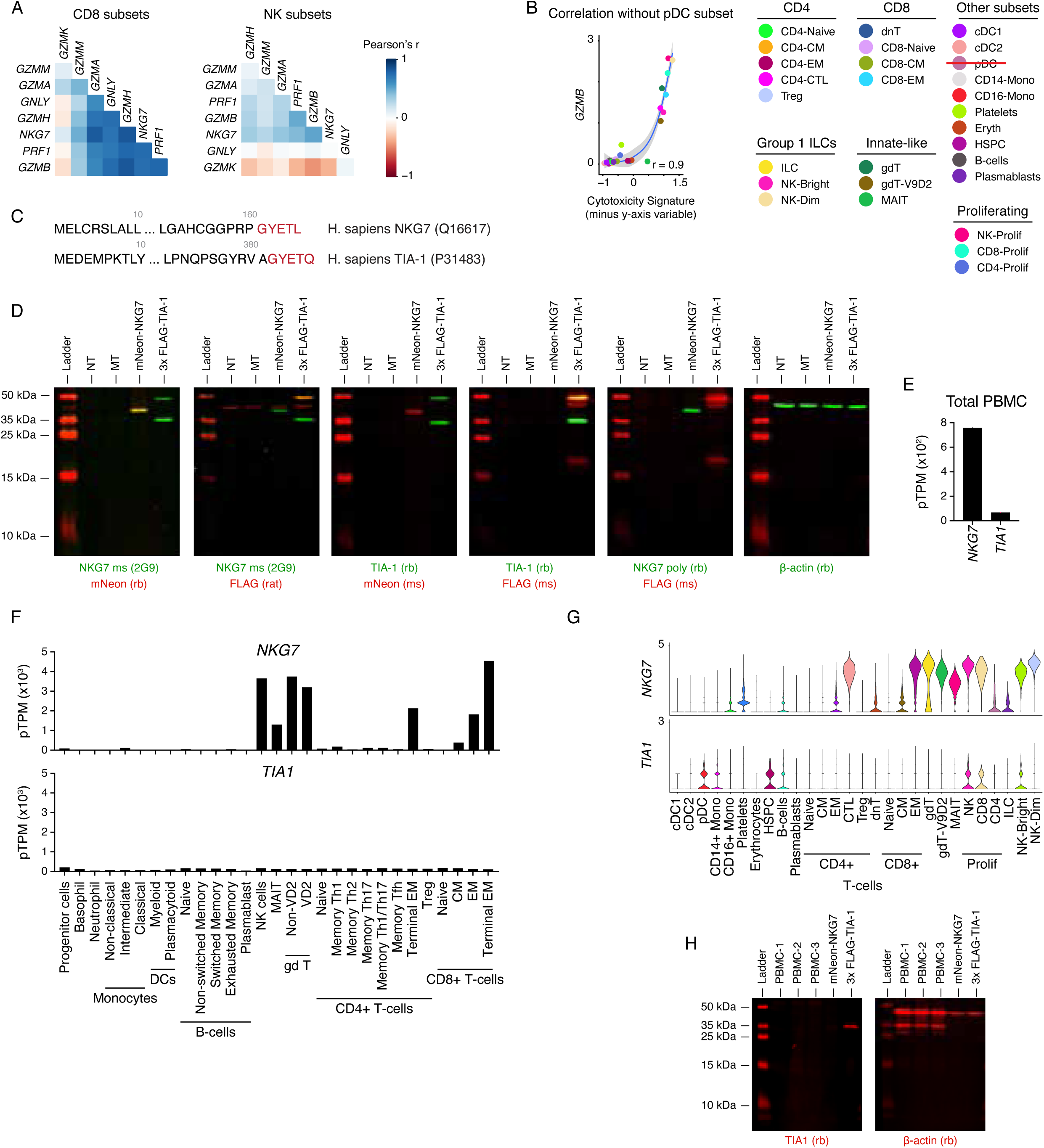
(A) Correlation of each cytotoxic gene with one another across populations in human PBMCs. (B) Scatterplots showing the correlation of *GZMB* with the cytotoxicity signature, excluding the Plasmacytoid Dendritic Cell (pDC) subset, in human PBMCs. Pearson correlation is displayed and the shaded area represents the 95% CI. (C) C-terminus sequences of TIA1 and NKG7, that are recognized by the 2G9 antibody. (D) Representative Western Blot showing that the 2G9 antibody is cross-reactive with TIA1 and NKG7. NT: non-transfected, MT: Mock-transfected. (E–G) Transcript levels of *NKG7* and *TIA1* in human PBMCs, in the whole population (E) and in the different sub-populations (F–G). Data in (E–F) are from bulk RNA-seq, while (G) represents single-cell RNA-seq. (H) Western Blot showing the absence of TIA1 in human PBMCs.

**Supplementary Figure 2.**
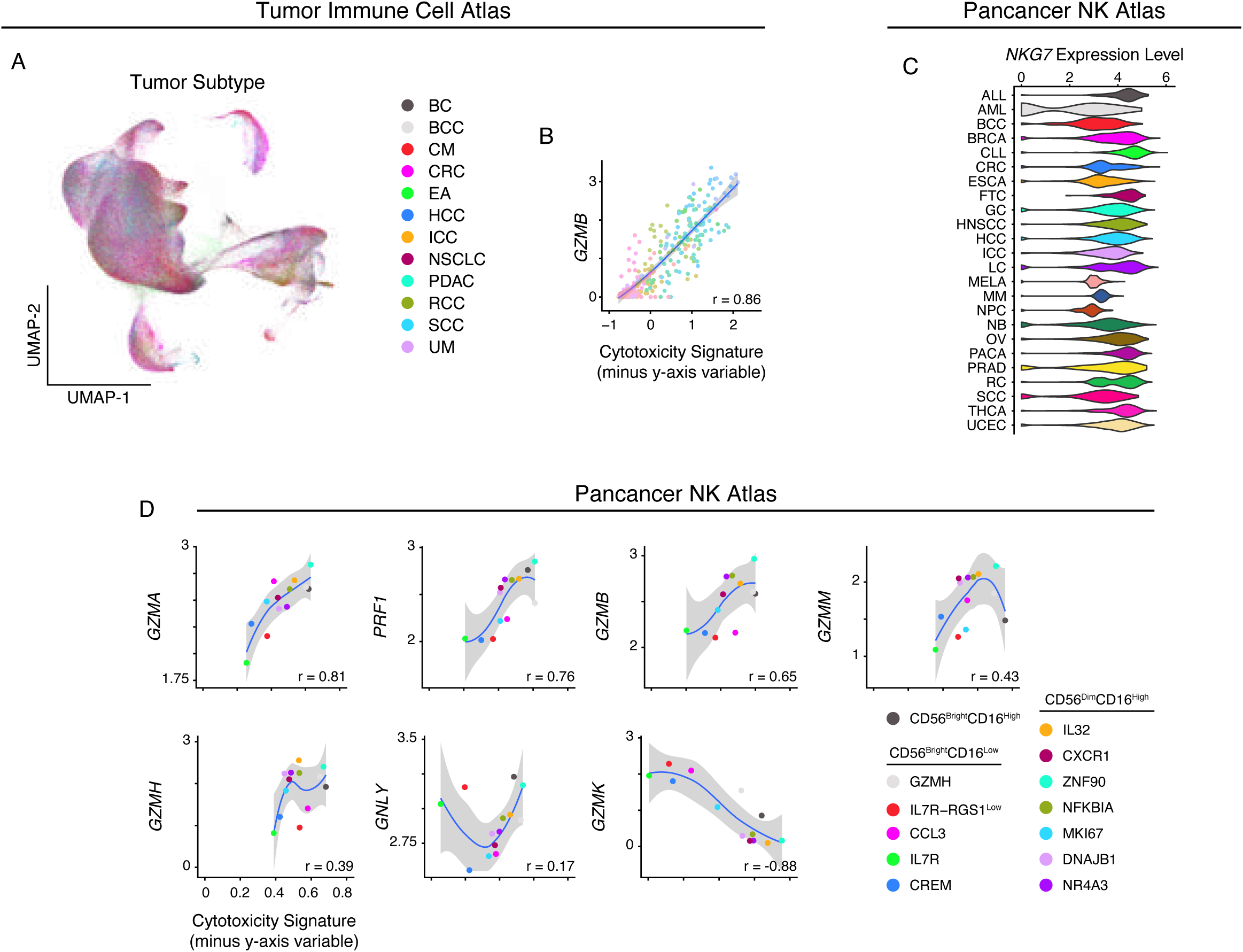
(A) UMAP plot of the different cell subsets identified within the tumor immune cell atlas. BC: Breast Cancer, BCC: Basal Cell Carcinoma, CM: Cutaneous Melanoma, CRC: Colorectal Cancer, EA: Esophageal Adenocarcinoma, HCC: Hepatocellular Carcinoma, ICC: Intrahepatic Cholangiocarcinoma, NSCLC: Non-small Cell Lung Cancer, PDAC: Pancreatic Ductal Adenocarcinoma, RCC: Renal Cell Carcinoma, SCC: Squamous Cell Carcinoma, UM: Uveal Melanoma. (B) Scatterplots showing the correlation of *GZMB* with cytotoxicity signature, in the cell subsets identified within the tumor immune cell atlas. Pearson correlation is displayed and the shaded area represents the 95% CI. (C) Violin pots showing the expression of *NKG7* in NK cells, as identified in the Pancancer NK Atlas. ALL: Acute Lymphocytic Leukemia, AML: Acute Myeloid Leukemia, BCC: Basal Cell Carcinoma, BRCA: Brest Cancer, CLL: Chronic Lymphocytic Leukemia, CRC: Colorectal Cancer, ESCA: Esophageal Cancer, FTC: Fallopian Tube Carcinoma, GC: Gastric Cancer, HNSCC: Head and Neck Squamous Cell Carcinoma, HCC: Hepatocellular Carcinoma, ICC: Intrahepatic cholangiocarcinoma, LC: Lung Cancer, MELA: Melanoma, MM: Multiple Myeloma, NPC: Nasopharyngeal Carcinoma, NB: Neuroblastoma, OV: Ovarian Cancer, PACA: Pancreatic Cancer, PRAD: Prostate Cancer, RC: Renal Carcinoma, SCC: Squamous Cell Carcinoma, THCA: Thyroid Carcinoma, UCEC: Uterine Corpus Endometrial Carcinoma. (D) Scatterplots showing the correlation of *GZMA*, *PRF1*, *GZMB*, *GZMM*, *GZMH*, *GNLY*, *GZMK* with the cytotoxicity signature, in NK cells, as identified within the Pancancer NK Atlas. Pearson correlation is displayed and the shaded area represents the 95% CI.

